# Endoglucanase-2 (Eng2), a conserved immunodominant antigen in dimorphic fungi that elicits immunity and resistance during infection

**DOI:** 10.1101/2025.06.23.661097

**Authors:** Uju Joy Okaa, Cleison Ledesma Taira, Lucas dos Santos Dias, Hannah Dobson, Gregory Kujoth, Althea Campuzano, Jane Homan, George R. Thompson, ChiungYu Hung, George S. Deepe, Marcel Wüthrich, Bruce S. Klein

**Affiliations:** Departments of Pediatrics, University of Wisconsin School of Medicine and Public Health, University of Wisconsin-Madison, Madison WI, USA; Internal Medicine, University of Wisconsin School of Medicine and Public Health, University of Wisconsin-Madison, Madison WI, USA; Medical Microbiology and Immunology University of Wisconsin School of Medicine and Public Health, University of Wisconsin-Madison, Madison WI, USA; Fungal Pathogenesis Section, Laboratory of Clinical Immunology & Microbiology (LCIM), National Institute of Allergy & Infectious Diseases, National Institutes of Health (NIH), Bethesda, MD; Department of Molecular Microbiology and Immunology, The University of Texas at San Antonio, San Antonio, TX 78249, USA; ioGenetics LLC, Madison, WI, USA; Department of Internal Medicine, University of California Davis Medical Center, Sacramento, CA 96817, USA; Department of Medicine, Division of Infectious Diseases, University of Cincinnati College of Medicine, Cincinnati,OH, USA

## Abstract

Herein, we describe a conserved surface and cell wall protein, Endoglucanase 2 (Eng2), expressed on the etiological agents that cause the endemic systemic mycoses of North America – *Blastomyces, Coccidioides and Histoplasma*. We demonstrate that despite sequence variation of the protein across these related fungi, exposure to Eng2 vaccinates and protects inbred and humanized HLA-DR4 strains of mice against lethal experimental infections with these fungi by eliciting adaptive immunity mediated by CD4 T cells. We also show that CD4 T cell precursors against Eng2 are detectable in naïve individuals and that patients who have recovered from these infections evince a memory and recall CD4 T cell response to Eng2 and its immunodominant epitopes that we have mapped. We create and catalogue new tools and information such as immunodominant peptide epitopes of Eng2 from each fungus recognized by inbred mice and human subjects and we engineer novel peptide-MHC II tetramers for tracking T cells in inbred and HLA-DR4 humanized mice that will be useful for those who study these infections in mice and humans. Lastly, because most patients demonstrate memory and recall responses against Eng2, our work offers new tools for diagnosis of this collection of infectious diseases across North America.

## INTRODUCTION

The endemic dimorphic fungi of North America including *Blastomyces, Coccidioides* and *Histoplasma* collectively cause over a million new cases each year and represent the most common cause of fungal pneumonia in otherwise healthy individuals. Many of these infections are mild or asymptomatic, but increasing numbers represent reactivation or new acquisition of severe disease in the setting of biologics and other forms of immune suppression. The endemic zones of these pathogens also are spreading likely due to anthropurgic behavior and climate.

Vaccination against infectious diseases has significantly improved public health. Once common and deadly bacterial and viral diseases are now rare or eliminated in vaccinated populations (1). Despite these successes and the growing clinical need, there are currently no fungal vaccines licensed for human use (2). A major limitation for developing vaccines against fungi stems from the lack of knowledge of highly protective antigens, especially those that are conserved and can protect against multiple fungi. Such immunodominant antigens also could be used for developing better methods of diagnosis.

We recently discovered *Blastomyces* endoglucanase 2 (Eng2) (or *Bl*-Eng2), a glycoprotein in *Blastomyces dermatitidis (Bd)* that harbors potent antigenic (3) and adjuvant properties (4). *Bl*-Eng2 elicits cellular immunity and protects against experimental infection with this fungus. A peptide-MHC tetramer specific to an immunodominant region of the antigen demonstrated that vaccination and infection expanded and recruited several hundred thousand T cells to the lung after infection (3). To our knowledge, this is the largest activation and expansion of functional anti-fungal T cells reported to date. The potency and functional versatility of this antigen and its conservation across the dimorphic fungi prompted us to investigate whether vaccination with *Bl*-Eng2 or its homologues could protect against infection with the other dimorphic fungi of North America and thus serve as a pan-fungal antigen for vaccination and diagnostics.

Herein, we report that vaccination with Eng2 homologues protects against experimental infection with corresponding dimorphic fungi in inbred C57BL/6 mice and humanized HLA-DR4 mice. We also show that CD4^+^ T cell precursors against these antigens are detectable in the blood of naïve human subjects and observe that patients who have recovered from each of these infections demonstrate a memory or recall CD4^+^ T cell response to the antigen. Given the immunological relevance of Eng2, we mapped the immunogenic T cell epitopes. Importantly, we now create and catalogue a compendium of new epitope information and related immunological tools such as the immunodominant peptides of the Eng2 homologue in each dimorphic fungus for both strains of mice and human subjects, as well as novel peptide-MHC II tetramers for tracking CD4^+^ T cells in inbred C57B/L6 and HLA-DR4 humanized mice. This new information and the accompanying tools will be invaluable for large numbers of investigators who study basic and clinical questions about immunity to these infections in mice and humans. Lastly, as most patients demonstrate memory immunity and recall responses against Eng2, our work offers new tools for diagnosis of this collection of important infectious diseases across North America

## RESULTS

### Eng2 homologues are conserved and associated with the cell wall of dimorphic fungi

Subcutaneous (SC) vaccination with *Bl*-Eng2 protects mice against experimental infection with *Bd* (3). In addition, we found that the amino acid sequences for the respective Eng2 homologues of the three dimorphic fungi *Bd*, *Coccidioides posadasii (Cp)* and *Histoplasma capsulatum (Hc)* are highly conserved (**Fig. 1A**). The sequence similarity for the three homologues ranges from 50% to 68% and the identity ranges from 38% to 60% (**Fig. 1B**). Given these results, we sought to test whether vaccination with *Bl*-Eng2 also protects mice against infection with *Cp* and *Hc*. Lung colony forming units (CFU) of *Bl*-Eng2 vaccinated mice were similar to unvaccinated mice after infection with *Cp* or *Hc* (**SFig. 1**). Thus, despite the sequence similarity and identity of Eng2 across these dimorphic fungi, vaccination with *Bl*-Eng2 does not cross-protect against experimental infection with *Cp* and *Hc*. We thus cloned the homologues of the respective dimorphic fungi, expressed them in *Pichia pastoris* (**Fig. 1C**), and raised antibodies against each protein homologue. Staining of yeast with these antibodies and analysis by fluorescent microscopy and FACS localized Eng2 homologues to the cell wall (**Fig. 1D+E**).

**Figure 1.**
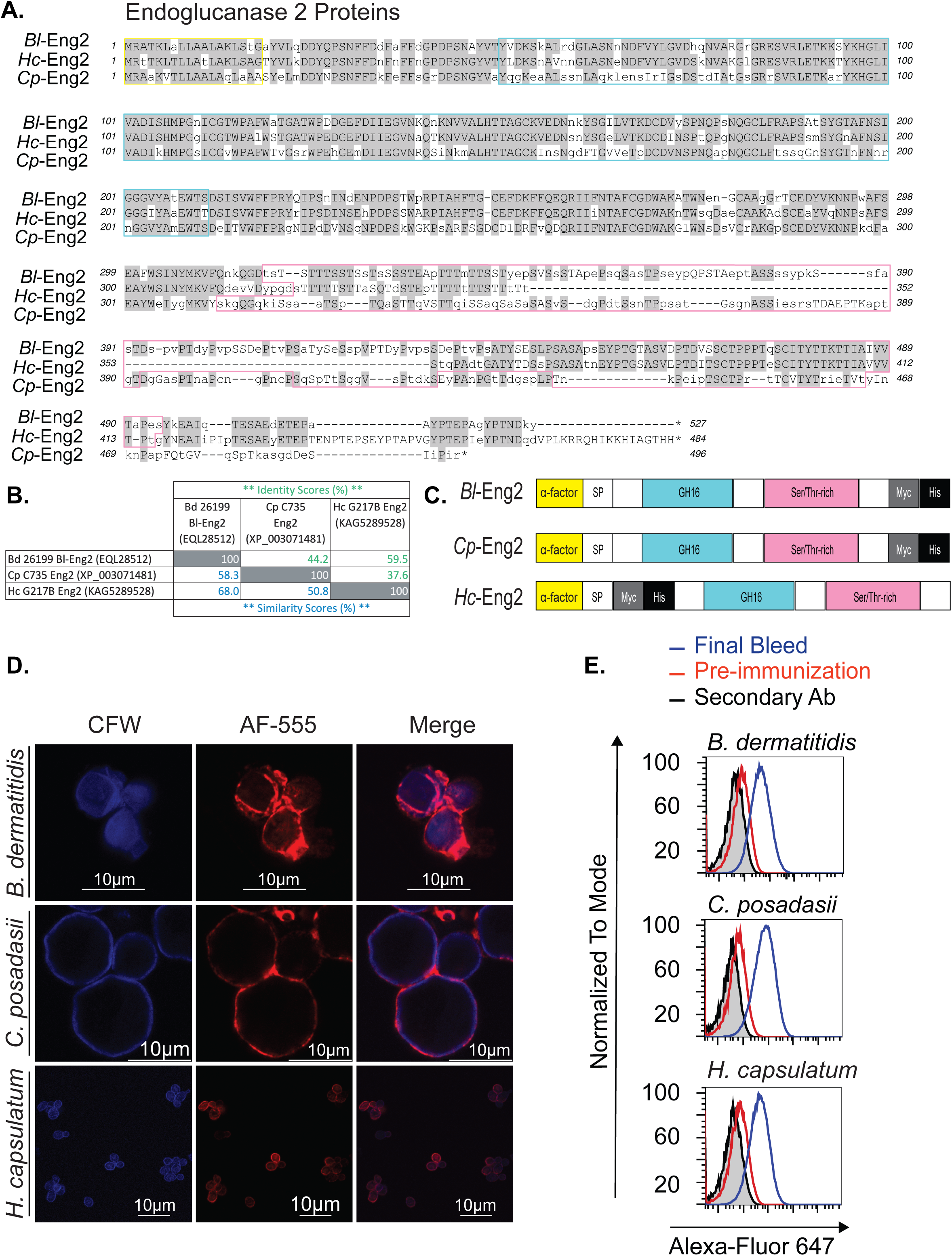
Alignment of Endoglucanase 2 amino acid sequences of dimorphic fungi. **A)** Amino acid sequence of endoglucanase 2 homologs (Eng2) (*Bl*-Eng2 = *Blastomyces dermatitidis* Eng2; *Cp*-Eng2 = *Coccidioides posadasii* Eng2; *Hc*-Eng2 = *Histoplasma capsulatum* Eng2). Colors represent domains denoted in panel C. **B)** The percentage of identical and similar amino acids between Eng2 homologues was determined by T-Coffee multiple sequence alignment. **C)** Domains of Eng2: yellow, α-factor secretory signal; SP, native Eng2 signal peptide; blue, GH16 domain; pink, serine/threonine-rich domain; gray, c-Myc-tag; and black, histidine tag. **D)** Immunofluorescence microscopy showing cell wall localization of Eng2 in *Bd* yeast, *Cp* spherules, and *H*c yeast. Calcofluor White Stain (CFW) of fungal chitin; AF (Alexa Fluor)-555 = Rabbit polyclonal α-Eng2 raised against each homologue. **E)** Flow cytometry analysis of *Bd* and *Hc yeast*, and *Cp* spherules, stained with rabbit pre-immune control serum and α-Eng2 rabbit immune serum.

### Protective efficacy of Eng-2 homologues against endemic dimorphic fungi

To investigate whether vaccination with *Cp*-Eng2 and *Hc*-Eng2 protects mice against respective infection with *Cp* and *Hc*, we purified recombinant proteins and generally loaded them into glucan-chitin particles (GCP) as described (5). Vaccination with *Cp*-Eng2 followed by infection with *Cp* reduced CFU in the lung and spleen by 3-4 logs compared to unvaccinated controls (**Fig. 2 A**). As a consequence of increased resistance, vaccinated mice did not lose weight as opposed to unvaccinated mice (**Fig. 2B**) and showed an increased survival rate following lethal pulmonary infection compared to unvaccinated controls (**Fig. 2C**). Similarly, vaccination with *Hc*-Eng2 protected C57BL6 mice against *Hc* following lethal pulmonary infection (**Fig. 2D-F**).

**Figure 2:**
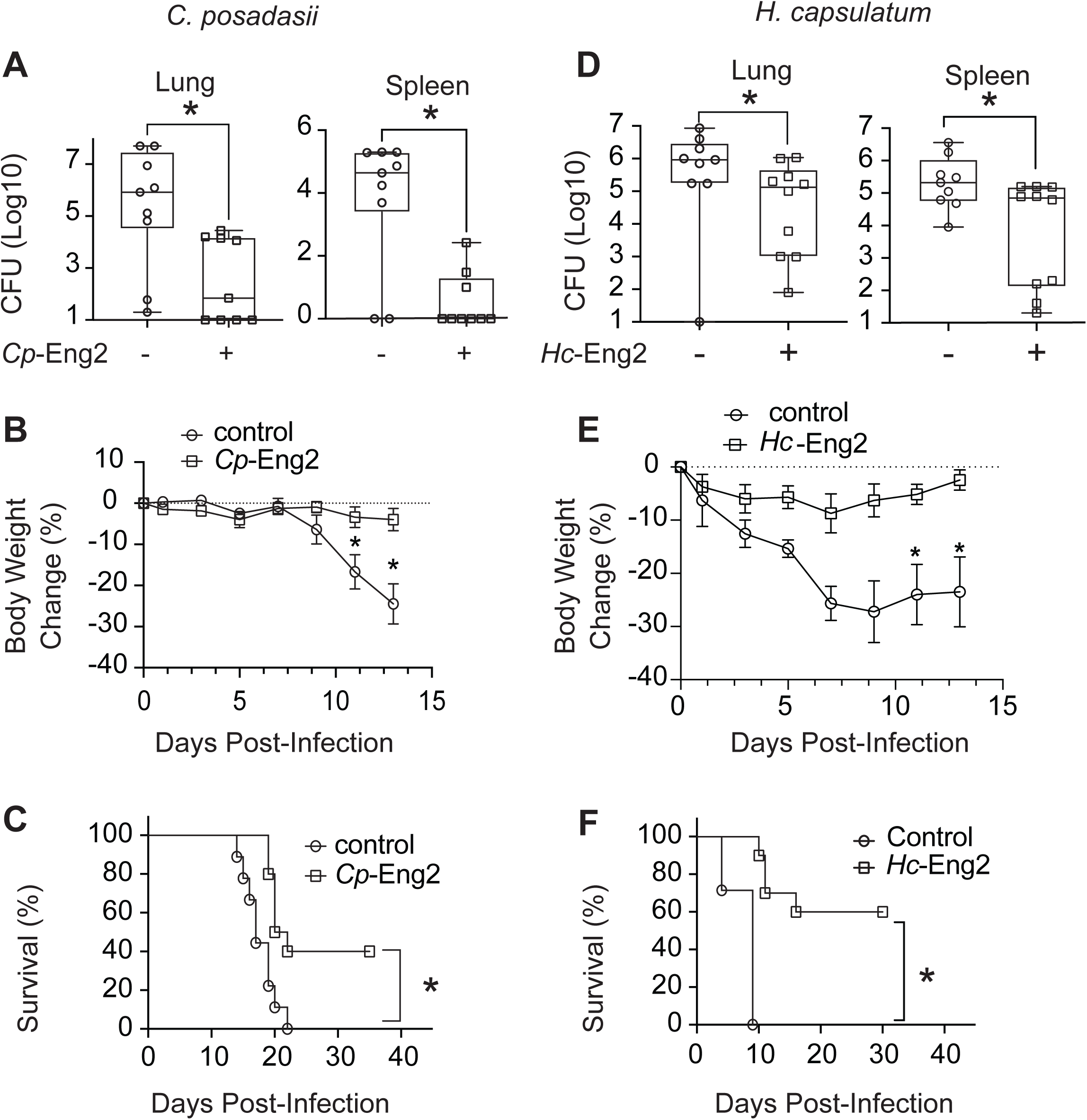
Protective eaicacy of Endoglucase-2 (Eng-2) against endemic dimorphic fungi. Homologous Eng2 proteins, *Cp*-Eng2 and *Hc*-Eng2 were respectively used to vaccinate C57BL/6 mice against of *Cp* (**A, B & C**) and *Hc* (**D, E & F**) as described in Methods. Burden of infection (colony forming units (CFU), weight loss, and survival are depicted (from top to bottom). Percent body weight change is depicted in mice monitored daily. Data shown are from a representative experiment of two to three performed (n=10 mice/group). CFU are expressed as Log_10_ plotted with geometric mean ± geometric SD. *p<0.05, Two tailed Mann-Whitney T test.

### Identification of immunogenic CD4^+^ T cell epitopes of *Cp*-Eng2 and *Hc*-Eng2 in C57BL6 mice

Since vaccine protection required immunization with homologous Eng2 proteins, we hypothesized that the immunogenic T cell epitopes are different for each homologue. To test this idea, we mapped the Eng2 peptide epitopes recognized by CD4^+^ T cells and generated class II MHC-peptide tetramers as described (5, 6). We analyzed the *Cp*-Eng2 and *Hc*-Eng2 homologues for MHC class II peptide-binding sequences. Of seven predicted peptides epitopes from these homologue, three 13-mers from each one triggered IFN-γ production by CD4^+^ T cells isolated from the spleen of mice vaccinated with the corresponding Eng2 protein (**Fig. 3A+D**). Peptide 3 from *Cp*-Eng2 (TVWFFPRGNIPDD) and peptides 1 and 5 from *Hc*-Eng2 (NFFNGPDPSNGYV and SSWARPIAHFTGC) induced the strongest IFN-γ response (**Fig. 3A, D+G**). Thus, mapping the position of the immunogenic *Cp*-Eng2 and *Hc*-Eng2 peptides indicated different locations of the immunodominant CD4^+^ T cell epitopes in each homologue (**Fig. 3G**). These regions also differed from that in *Bl*-Eng2 (data not shown) (3).

**Figure 3:**
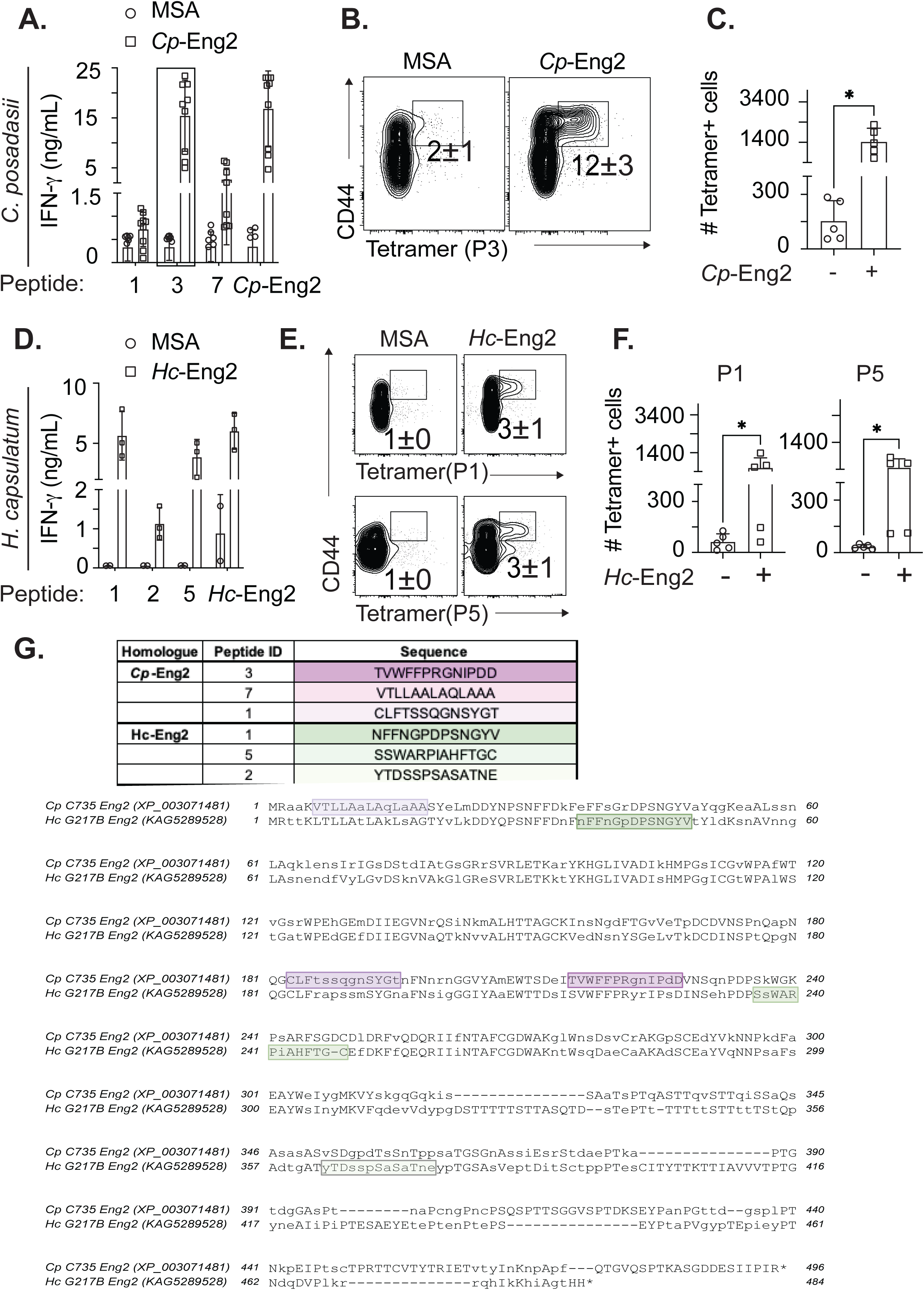
Mapping of Eng2 immunodominant epitopes in *C. posadasii* and *H. capsulatum* in C57BL/6 mice. **A+D)** Identification of immunodominant epitopes of *Cp*-Eng2 and *Hc*-Eng2. Mice (n=5/group) were vaccinated with *Cp*-Eng2 and *Hc*-Eng2 as described in Methods. Two weeks after the boost, splenocytes were restimulated *ex vivo* with peptide P3 from *Cp*-Eng2 (**C & D**) or P1 and P5 from *Hc*-Eng2. (**B,E,C,F)** Splenocytes from vaccinated mice were stained with tetramers containing peptide P3 (*Cp*-Eng2), P1 or P5 (*Hc*-Eng2) and analyzed by flow cytometry. The frequency and numbers of tetramer^+^ cells are illustrated in panels (**B,E,C,F**). The data shown are representative of two independent experiments. Data are represented as mean± SEM and analyzed using the T-test and Mann-Whitney U test. *p=<0.05. (**G)** *Cp*-Eng2 and *Hc*-Eng2 sequences were aligned with the T-coffee algorithm using MacVector 18.5.1. Experimentally determined immunodominant epitopes are highlighted in purple and green gradient-shaded boxes for *Cp*-Eng2 and *Hc*-Eng2, respectively. Peptides that elicited the most IFN-γ (*Cp*-P3 and *Hc*-P1) are shaded dark purple or green.

We used the immunodominant peptides to create MHC class II tetramers and used them to demonstrate expansion of primed Eng2 antigen-specific, memory CD4^+^ T cells in the spleens of vaccinated mice (**Fig. 3B+C and E+F**). About 12% of CD4^+^ T cells stained with the tetramer and the CD44 activation marker after vaccination with *Cp*-Eng2 and about 3% of CD4^+^ T cells did so following vaccination with *Hc-Eng2*. We validated that tetramer binding was specific since few CD8^+^ T cells bound the respective tetramers (**SFig. 2A).** The precursor frequency of naïve (CD44^low^) Eng2-specific CD4^+^ T cells in unvaccinated mice was 32 for *Hc*-Eng2, 97 for *Bl*-Eng2 and 130 for*Cp*-Eng2) (**SFig. 2C**). Vaccination with corresponding Eng2-homologues activated and expanded tetramer^+^ CD4^+^ T cells by 7- to 29-fold compared to the numbers present in naïve mice (**SFig. 2B+C**). In sum, mapping the position of the immunogenic *Cp*-Eng2 and *Hc*-Eng2 peptides indicated different locations of the T cell epitopes (**Fig. 3G**). Thus, we have generated tetramers against all three homologues of Eng2 proteins, which allow tracking and enumeration of Eng2-specific CD4^+^ T cells during the evolution of immunity against these endemic systemic mycoses.

### Functional analysis of Eng-2-specific CD4^+^ T cells in vaccinated C57BL/6 mice following infection with *Cp* and *Hc*

We previously reported that *Bl*-Eng2 specific CD4^+^ T cells are recalled to the lung in substantial numbers upon infection in vaccinated C57BL/6 mice (3). We investigated here whether *Cp*- and *Hc*-Eng2-specific CD4^+^ T cells also accumulate in the lung in substantial numbers in vaccinated mice infected with the respective fungus. The frequencies and numbers of activated (CD44^+^), tetramer^+^ T cells were at least 10 times higher in vaccinated mice compared to unvaccinated mice at day 6 or 5 post-infection with *Cp* or *Hc, respectively* (**Fig. 4A+B and 4E+F**). Since vaccine immunity to dimorphic fungi is primarily mediated by Th1 and Th17 cells that respectively produce IFN-γ and IL-17 (5, 7–11), we stimulated primed T cells with the peptides that were used to generate the tetramers and measured cytokine production by intracellular staining. The frequencies and numbers of IFN-γ and IL-17 producing T cells from vaccinated mice was significantly increased compared to unvaccinated controls and indicated a mixed Th1/Th17 phenotype (**Fig. 4C+D and 4G+H**). Thus, the tetramer^+^ cells are functional with respect to cytokine production.

**Figure 4:**
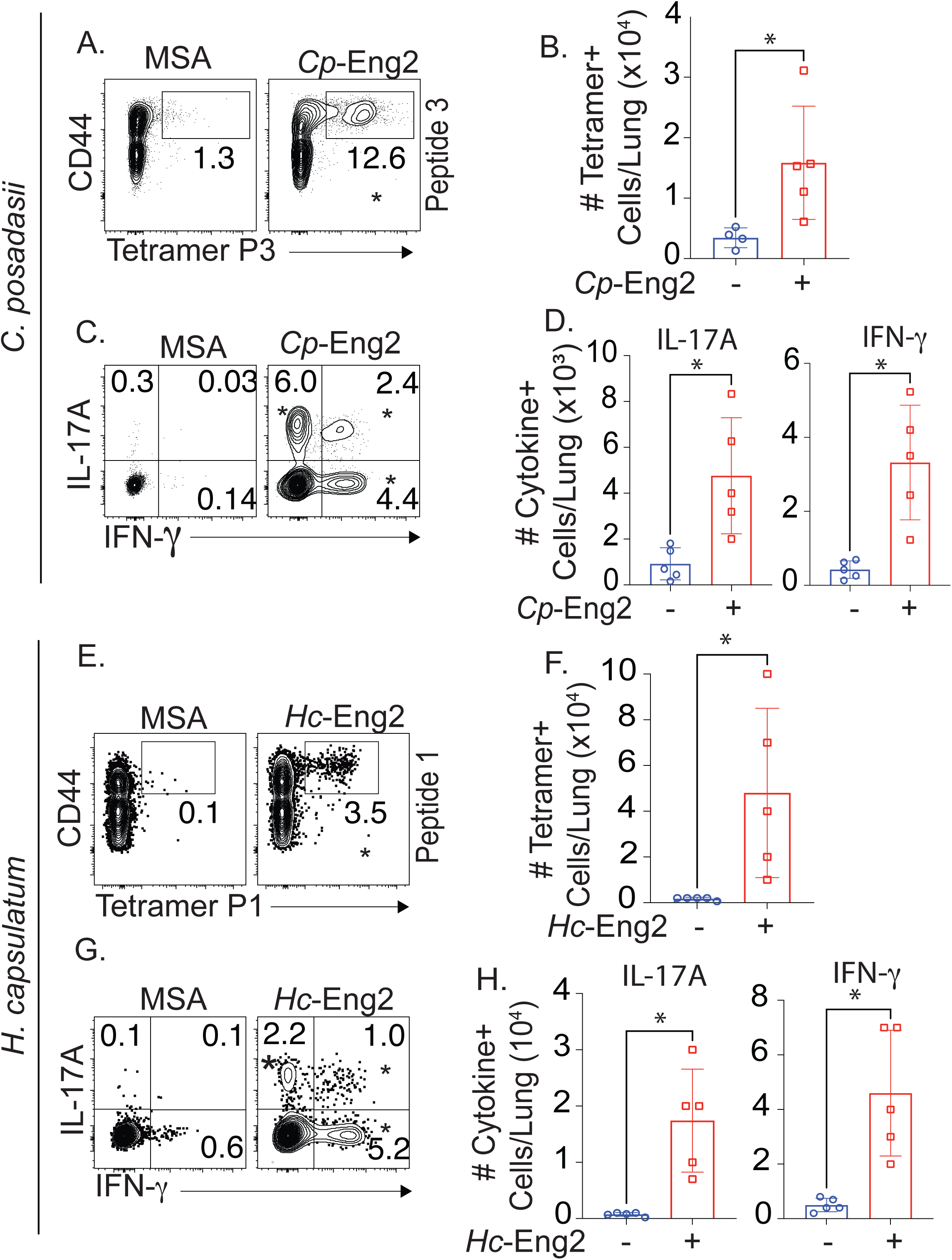
Functional analysis of Eng2 specific CD4^+^ T cells following experimental pulmonary infection with *C. posadasii* and *H. capsulatum*. Tetramer^+^ cells were determined after C57BL/6 mice were vaccinated as described in methods with *Cp*-Eng2 (**A-D**) or *Hc*-Eng2 (**E-H**) (n=5/group). Mice were experimentally infected as in methods and sacrificed 4 or 5 days post challenge followed by enumeration of CD4^+^ tetramer^+^ T cells in the lung. Panels A&E show contour flow plots with the percentages of tetramer^+^ CD4^+^ T cells; panels B&F show histogram bar graphs with the total number of tetramer^+^ cells. After intracellular staining of IFN-γ and IL-17, the frequencies (**C+E**) and numbers (**D+H**) of cytokine-producing cells were analyzed. Data were analyzed using Mann-Whitney U test *p<0.05 vs. control group.

### Vaccine protection conferred by immunodominant peptides vs. full-length Eng2 proteins in C57BL/6 mice

Vaccination with full-length recombinant proteins offers the advantage that multiple epitopes likely induce a polyclonal and multi-epitope driven T cell response that could result in optimal protection. Alternatively, some epitopes could activate Treg cells that dampen a vaccine benefit (12). Thus, identifying protective T cell epitopes and combining them in multi-epitope vaccines could provide augmented vaccine efficacy (13, 14). Here, we investigated CD4^+^ T cell responses and protective efficacy elicited by immunodominant peptides vs. full length recombinant Eng2 homologues. Vaccination with peptide 3 from *Cp*-Eng2 generally induced the expansion of tetramer^+^ CD4^+^ T cells, and the development of Th1 and Th17 cells (**Fig. 5A**). Similarly, peptides P1 and P5 from *Hc*-Eng2 and P1 from *Bl*-Eng2 each induced tetramer^+^ Th1 and Th17 cells that were comparable with the CD4^+^ T cell responses in mice vaccinated with the respective full-length protein (**Fig. 5B+C**). Although the tetramer response to vaccination with peptide 1 in *Cp*-Eng2 was lower than that for the full-length protein (**Fig. 5A**), peptide 1 reduced lung CFU of *Cp by* several logs, similar to that with the full-length protein (**Fig. 5D**). Likewise, vaccination with peptide 3 from *Bl*-Eng2 reduced lung CFU similar to that observed with the full-length protein. Conversely, vaccination with P1 and P5 from *Hc*-Eng2 combined did not reduce lung CFU (**Fig. 5D**). This is likely due to the overall modest protection achieved by the *Hc*-Eng2 protein. Nevertheless, peptide vaccine was as effective as full-length protein in protecting against *Cp* and *Bd*.

**Figure 5:**
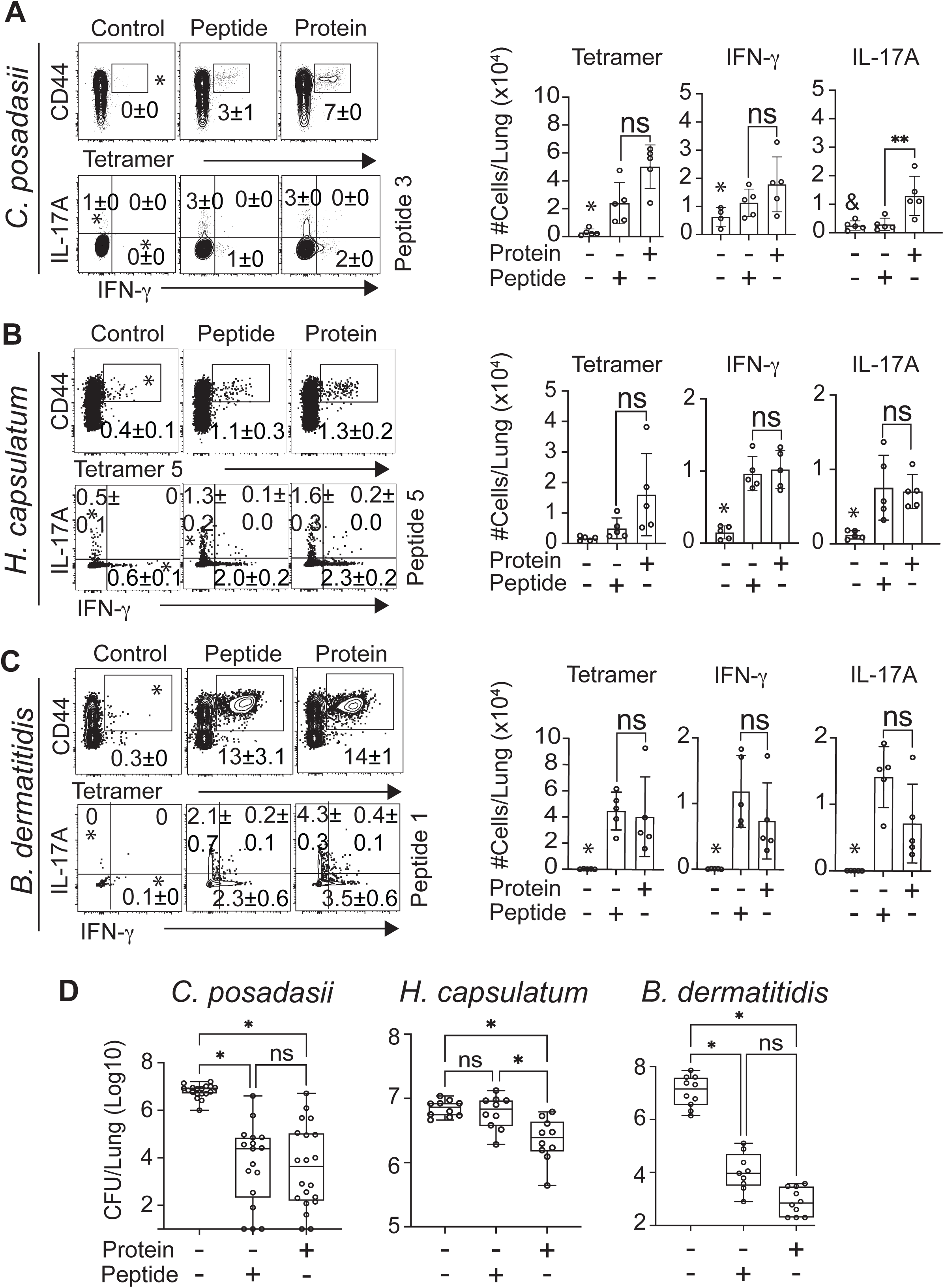
Vaccine protection conferred by immunodominant peptide vs. full-length protein homologue in wild type C57BL/6 mice. Mice were vaccinated with *Cp*-Eng2, *Hc*-Eng2 and *Bl*-Eng2 or equimolar amount of immunodominant peptides from each homologue as described in the Methods. Three weeks after the last boost, mice were challenged with a lethal dose of each organism. **(A-C)** Lung CD4^+^ T cells were analyzed for the frequencies and numbers of tetramer^+^ cytokine-producing T cells at day 6 (*Cp*), day 5 (*Hc*) and day 4 (*Bd*) post-infection (n= 5 mice/group). **(D)** Lung CFU were analyzed two weeks post-infection when the control group was moribund (n=10 mice/group). CFU are expressed as Log_10_ plotted with geometric mean ± geometric SD. *p<0.05 vs. all other groups, ^&^p<0.05 vs. protein vaccinated. Data were analyzed using the T test and two tailed Mann-Whitney T test).

### Protective efficacy of Eng-2 homologues and mapping of T cell epitopes in humanized DR4 mice

To translate our findings from C57BL/6 mice toward humans, we vaccinated humanized HLA-DR4 (DRB1*0401) transgenic mice that express human MHC II and lack murine endogenous MHC Class II (15). Vaccination with *Cp*-Eng2, *Hc*-Eng2 or *Bl*-Eng2 each protected HLA-DR4 mice against infection with the corresponding fungus (**Fig. 6A**). HLA-DR4 mice vaccinated with *Cp*-Eng2 also lost less weight and survived longer than controls (**SFig. 3A+B**).

**Figure 6:**
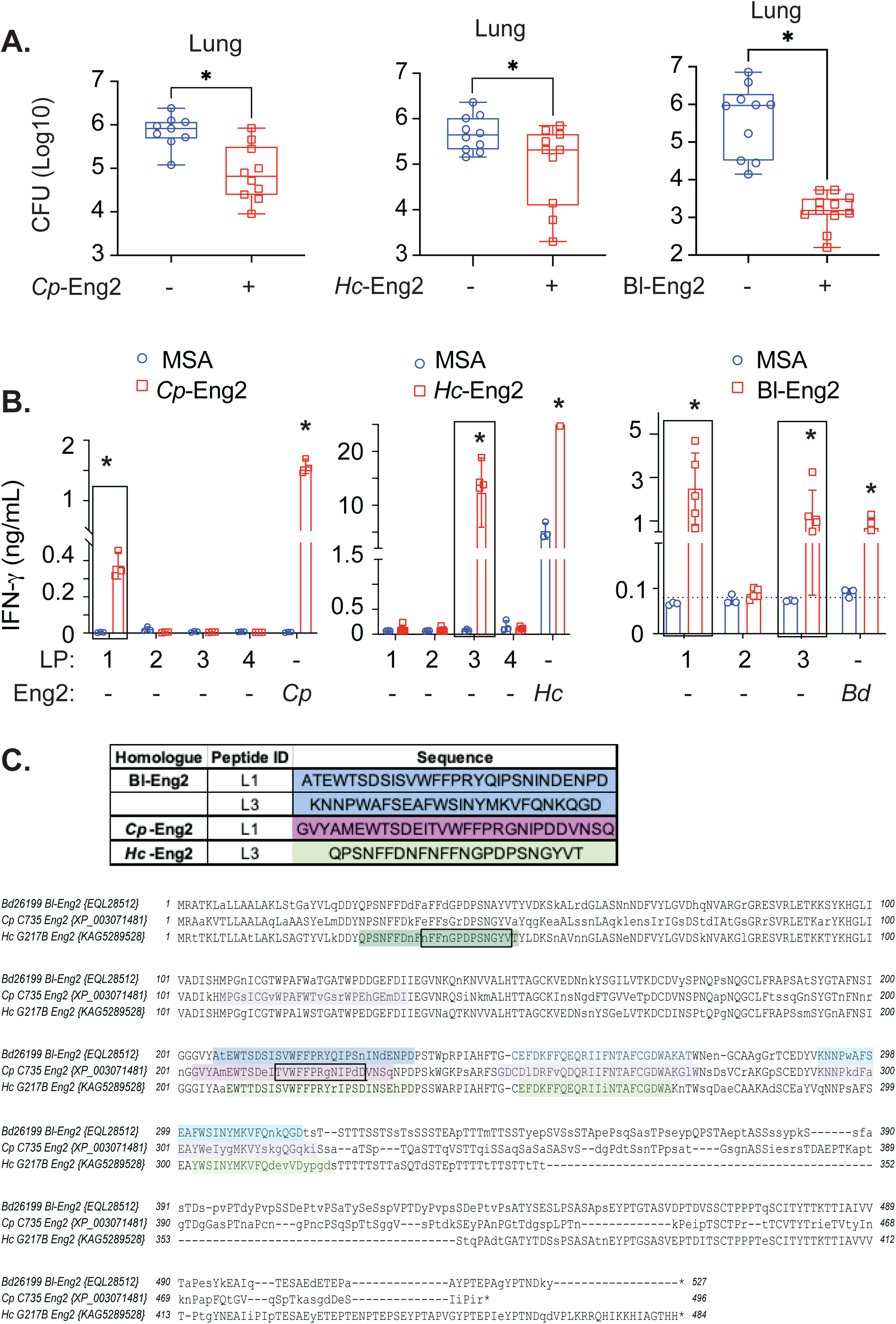
Protective eaicacy of Eng-2 homologues and mapping of Eng2 epitopes in humanized HLA-DR4 mice. **(A)** Mice were vaccinated with the respective Eng2 homologues and challenged and sacrificed as described in methods. CFU are expressed as Log_10_ plotted with geometric mean ± geometric standard error. (**B)** To assay peptide recognition, splenocytes from mice vaccinated with Eng2 homologues were stimulated *ex vivo* with predicted peptides that ranged from 25-30 amino acids. Cell culture supernatants were assayed for IFN-γ after five days of stimulation. *p<0.05 vs. corresponding MSA control groups. (**C)** The sequences of the Eng2 homologues were aligned with the T-coffee algorithm using MacVector 18.5.1. Experimentally determined immunodominant HLA-DR4 epitopes are highlighted in colors. Peptides that elicited the strongest IFN-γ response (*Cp*-P1, *Hc*-P3 and *Bd*-P1) are shaded in darker colors. The 13 mers of the immunodominant epitopes indicated in black boxes are shared between C57BL6 and HLA-DR4 mice.

To test whether a natural infection induces immunity to Eng-2, HLA-DR4 mice were sublethally infected with *H*c and their splenocytes were analyzed for cytokine responses to restimulation with *Hc*-Eng2 protein and peptides. CD4^+^ T cells primed during infection produced IFN-γ when restimulated *ex vivo* with *Hc*-Eng2 protein but not peptides (**SFig. 3F**). Thus, *Hc*-Eng2 is expressed on *Hc* yeast during pulmonary infection and *Hc*-Eng2-specific CD4^+^ T cells are induced during infection. Below, we demonstrate that PBMC from individuals that have recovered from *Hc* infection also react to *Hc*-Eng2 protein and peptides and therefore harbor antigen-specific memory CD4^+^ T cells.

We next determined the immunogenic peptides of Eng2 proteins that are recognized by human HLA-DR4. Using Immunoinformatics analytics software from Eigenbio™, we predicted peptides for each Eng2 homologue. We initially synthesized long peptides (LP) ranging between 25-30 amino acids to accommodate recognition of multiple HLA’s (in addition to DR4) in testing immunogenicity with human blood samples below. To identify immunogenic peptides, we recalled Eng2 homologue-primed T cells *ex vivo* with these peptides. Peptides LP1 from *Cp*-Eng2, LP3 from *Hc*-Eng2 and LP1 and LP3 from *Bl*-Eng2 triggered IFN-γ production (**Fig. 6B**). To map the nine core amino acids (plus 3 flanking aa on either side) that bind to HLA-DR4 and are recognized by the cognate T cell receptor (TCR), we synthesized 15-mers that overlap by 14 aa (referred to as peptide walking). GVYAMEWTSDEITVW (peptide 1) from *Cp*-LP1; NFNFFNGPDPSNGYV (peptide 1) from *Hc*-LP3; and PRYQIPSNINDENPD (peptide 15) from *Bd*-LP1 and VKNNPWAFSEAFWSI (peptide 16) from *Bd*-LP3 were identified as the immunogenic 15 mers (**SFig. 3C, E, G**). The sequence and location of the most immunogenic peptides in each of these Eng2 proteins restricted by HLA class II are depicted in **Fig. 6C**.

To validate our peptide mapping, we created a tetramer to track *Cp*-Eng2-specific CD4^+^ T cells in HLA-DR4 mice. After vaccinating mice with *Cp*-Eng2 and infecting them with *Cp*, we detected 4 % tetramer^+^ CD4^+^ T cells in the lung (**SFig. 3D**). We assessed vaccine immunity and protection by immunodominant peptides in HLA-DR4 mice. Vaccination induced the generation of IFN-γ producing CD4^+^ T cells (**SFig. 4A+C+E**). However, only vaccination with immunogenic peptides from *Cp*-Eng2 was able to reduce lung CFU in HLA-DR4 mice (**SFig. 4B+D+F**). We used the long immunogenic peptides identified in HLA-DR4 mice to test and confirm reactivity by naïve precursor CD4^+^ T cells in human subjects as outlined below.

### Identification of immunogenic peptides recognized by naïve CD4^+^ T cells from human subjects

To identify naïve human precursor CD4^+^ T cells that recognize Eng2 homologues and their immunogenic peptides, we exploited the Sanofi’s System as outlined in methods (**Fig.7A**). Briefly, antigens were loaded onto CD14^+^ monocytes and naïve human CD4^+^ T cells primed for 14 days. Primed T cells were restimulated with “fresh” dendritic cells (DCs) and stimulated with antigens for 5 to 7 hours while blocking intracellular protein transport to accumulate cytokines. CD4^+^ T cells from naïve donors produced increased intracellular IFN-γ, IL-2, TNF-α, IL-4 and IL-17 after Eng2 antigen priming and restimulation with the antigen compared to medium alone (**Fig. 7B**). The majority of primed CD4^+^ T cells were polyfunctional and produced multiple cytokines as indicated by Boolean analysis (**Fig. 7C**). The limited responses to *Hc*-Eng2 and peptides likely stemmed from the low precursor frequency (**Fig. S2C**). Interestingly, peptide primed CD4^+^ T cells showed higher stimulation indices for activated (CD154^+^) and IFN-γ producing cells than did the full-length protein-primed cells when restimulated with peptides (**Fig. 7D**). We assessed whether different immunodominant peptides from an individual Eng2 protein influenced the T helper phenotypes in an HLA haplotype-specific manner. Peptides 2 and 3 from *Cp*-Eng2 and peptide 1 from *Bl*-Eng2 favored the development of Th1 cells when presented by most HLA-DR haplotypes (**Fig. 7E**) although the sample size was insufficient to achieve statistical significance.

**Figure 7:**
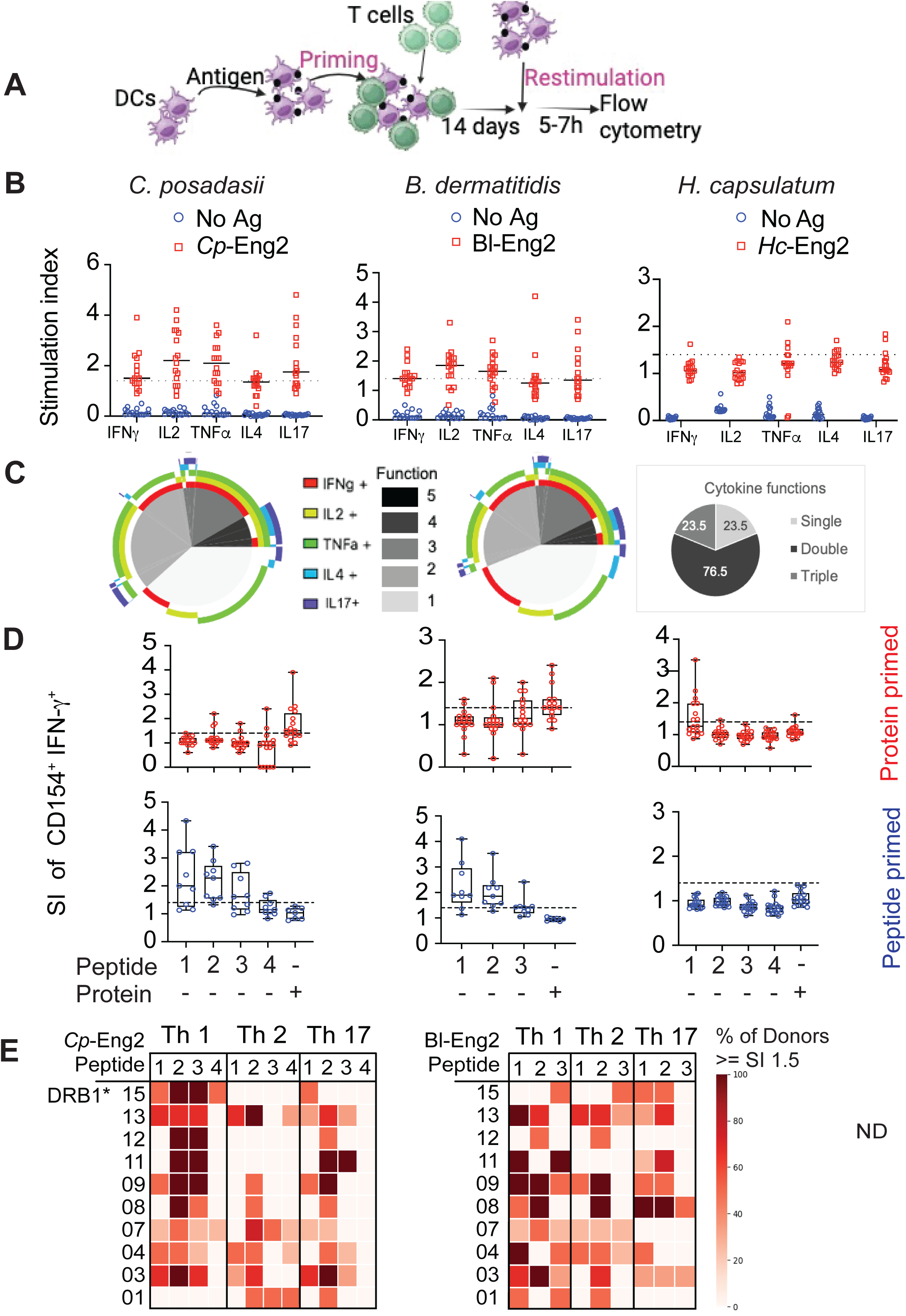
Response of Naïve human subjects to predicted epitopes. **(A)** Schematic of antigen priming strategy using Sanofi’s Modular Immune *in vitro* Construct (MIMIC) System as described in the Methods. **(B)** Naive CD4^+^ T cells from 16 healthy donors were primed and restimulated with full length Eng2 protein homologues from *Cp*, *Bd*, and *Hc* using the MIMIC System. Data are expressed as stimulation index (SI), which is the ratio of cytokines produced by cells that were primed and restimulated vs. cultured in medium alone. **(C)** Boolean analysis of the multifunctional cytokine response depicted in panel B, illustrating the fraction of activated donors CD4^+^ T cells that produced more than one cytokine following Eng2 stimulation (color); “function” denotes the number of cytokine products. (**D**) Response of naïve donor-derived CD4^+^ T cells to priming with protein (top row) or peptide pool (bottom row) and re-stimulation with individual peptides or protein in the MIMIC System. **(E)** Heat map of responses to peptides P1-P4 with different Th profiles according to individual HLA haplotype among the naïve study cohort.

### Memory T cell responses from patients recovered from infection with dimorphic fungi

We investigated patients who recovered from infection with *Cp*, *Hc* or *Bd* to assess whether they harbor a pool of memory CD4^+^ T cells specific for the corresponding Eng2 protein homologue and respective immunogenic peptides identified above. PBMCs from 12 patients who had recovered from confirmed infection with *Cp* all responded to positive control antigens (attenuated *Cp* strain *ΔCps1* (16–19) and *C. albicans*) and also produced IFN-γ and IL-17 in response to stimulation with *Cp*-Eng2 protein (**Fig. 8A**). These patients produced IFN-γ in response to *Cp*-Eng2 peptides 1-3 (**Fig. 8B**) as in the MIMIC System (**Fig. 7D**) but only produced IL-17 in response to peptide 3 in this assay (**Fig. 8B**).

**Figure 8.**
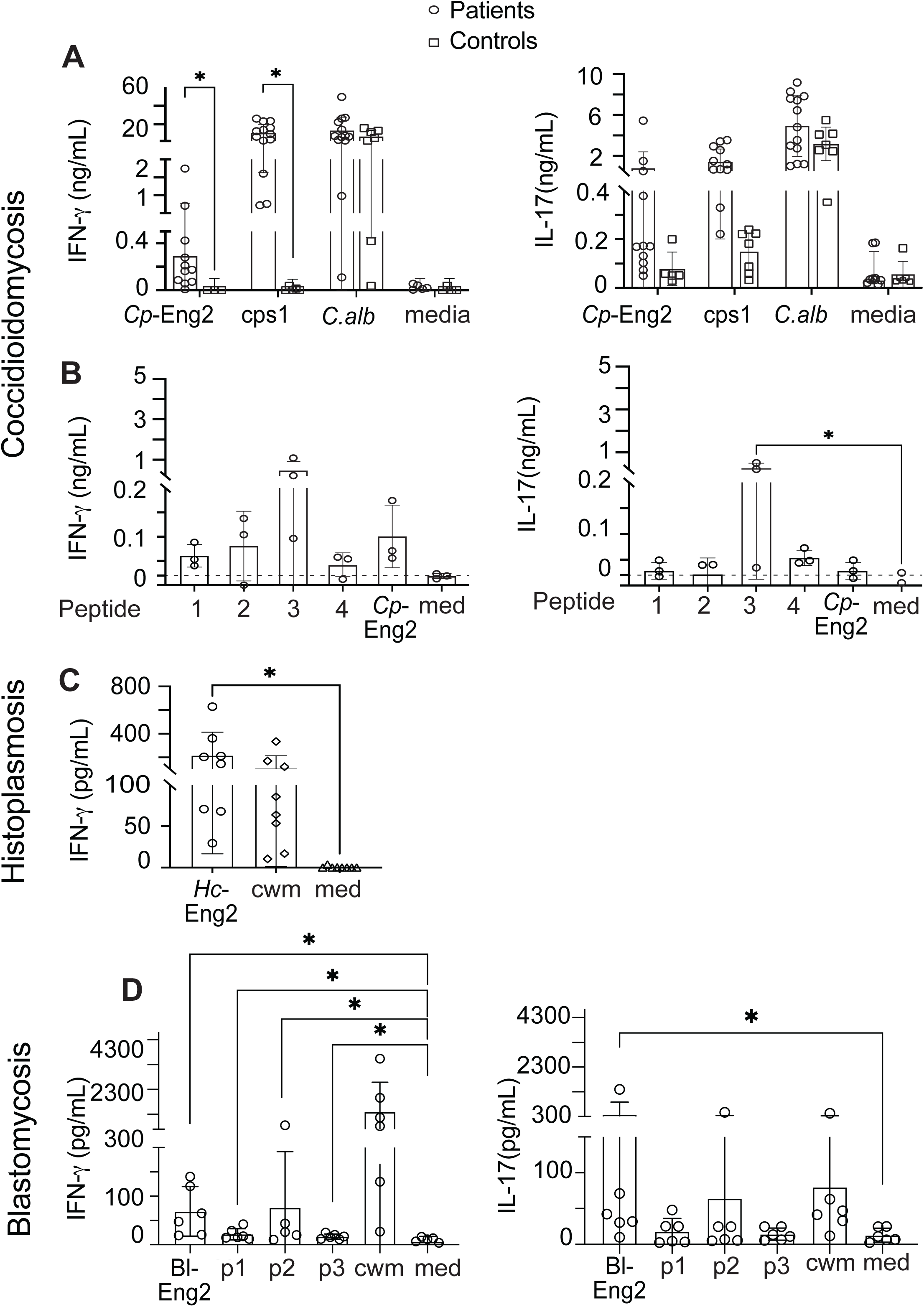
Eng2-specific memory T cells in fungi exposed human subjects. Recall response of CD4^+^ T cells from patients with coccidioidomycosis and healthy control subjects after stimulation with (**A**) *Cp*-Eng2 protein or (**B**) peptides. CD4^+^ T cells were expanded by co-culturing them with CD14^+^ monocytes in the presence of various stimuli for 7 days. IFN-γ and IL-17 were measured in the culture supernatant by ELISA. Heat killed *11cps1* spores were used to confirm the patient harbored immunity to *Cp* and *Candida* yeast served as a control to exclude anergy. Stimulation media alone served as negative control. (**C**) *Ex-vivo* response of PBMC from healthy blood donors previously exposed to *H. capsulatum* following 5 days of stimulation with *Hc*-Eng2 or *Hc* CWM antigen as an indicator of prior infection. Each symbol represents a patient or control. Data was analyzed using Mann-Whitney U test. *p=<0.05.

About 80% of people who live in the *Hc* endemic region of Ohio have evidence of prior infection with the fungus. Such persons were identified by production of IFN-γ by their PBMCs in response to a positive control stimulus of cell wall membrane (CW/M) extract from *Hc.* Eight of 10 individuals had evidence of CW/M reactivity and also responded strongly to *Hc*-Eng2 (**Fig. 8C**). None of the subjects that failed to respond to CW/M responded to *Hc*-Eng2, supporting the specificity of the response to *Hc*-Eng2.

PBMCs from all six patients who had recovered from culture-confirmed infection with *Bd* and responded to the positive control stimulus of CW/M extract from *Bd* also responded to *Bl*-Eng2 (**Fig. 8D**). Their cells also produced IFN-γ and IL-17 following stimulation with *Bl*-Eng2 protein and peptides 1-3. Two other confirmed cases with acute infection in whom treatment was just initiated failed to respond to a positive control stimulus and were unevaluable.

From these results, we conclude that the majority of subjects who had recovered from infections with each of the endemic mycoses responded to the corresponding Eng2 homologue and also to the immunogenic peptides discerned above in studies of humanized mice and naïve human subjects.

## DISCUSSION

The endemic systemic mycoses are invasive fungal diseases that usually occur in previously healthy individuals and are not limited to hosts with impaired cellular or humoral immunity. We report here that Eng2 from the cell wall of the causative fungi elicits immunity in both humans and mice and protects the latter against experimental lethal pulmonary infection. Thus, vaccination with the respective Eng2 homologue protected C57BL6 and humanized HLA-DR4 mice against corresponding infections with *Cp*, *Hc* and *Bd*. Unexpectedly, despite the high level of sequence conservation among the Eng2 homologues, we did not observe cross-protection by *Bl*-Eng2 against infection with *Cp* and *Hc*. Nevertheless, each of the respective homologues induced resistance in mice and adaptive immunity in humans during their infections. Thus, persons recovered from infection displayed reactivity to the respective protein suggesting that these antigens could have diagnostic value for detection of infection.

By mapping immunodominant T cell epitopes for each Eng2 homologue, we learned that the peptides differed in sequence and location. This contrasts with our findings with another conserved antigen, calnexin, that conferred vaccine protection against *Cp*, *Hc* and *Bd* by an identical 13-amino acid epitope (5). Corresponding 1807 T cell receptor transgenic (calnexin-specific) T cells expanded not only after infection with *Bd* but also after challenge with other dimorphic fungi including *Cp* and *Hc* (20). Furthermore, infection with other clinically relevant ascomycetes such as *Aspergillus fumigatus*, *Fonsecaea pedrosoi* and *Pseudogymnoascus destructans* triggered the expansion of calnexin peptide-MHC II tetramers (5). The calnexin study thus demonstrated the possibility of exploiting a single conserved epitope/antigen as a target for the development of broadly reactive antifungal vaccines. However, a limitation for the development of a calnexin-specific antifungal vaccine is the low precursor frequency of naïve endogenous CD4^+^ T cells and the modest expansion and protective efficacy after vaccination and infection (5). We overcame this limitation here with the discovery of Eng2 homologues. The precursor frequencies for the immunodominant epitopes of *Cp*- and *Bl*-Eng2-specific naïve endogenous CD4^+^ T cells in C57BL/6 mice is 97 to 130 cells, compared to only 29 for calnexin-specific (5) and 32 to 47 for *Hc*-Eng2-specific precursors. Consequently, vaccine-induced expansion of antigen-specific T cells and protection was much higher for experimental *Cp-* and *Bd-*infection compared to that for *Hc*-Eng2. Our data support the concept that a large population of T cell precursors is associated with increased TCR diversity and larger primary immune response and explains why immune responses to some peptides are stronger than others (21).

We translated our findings from C57BL6 mice to humans. We first vaccinated humanized HLA-DR4 mice to confirm protection and identify the immunogenic T cell epitopes. Vaccination with all three Eng2 homologues protected HLA-DR4 mice against infection with the respective dimorphic fungi, suggesting that human T cells could be primed during immunization. This idea was supported by the observation that vaccination with Eng2 homologues and peptides primed HLA-DR4 restricted CD4^+^ T cells that produced IFN-γ upon recall *in vitro*. Since the immunogenic epitopes for the three Eng2 homologous differed in amino acid sequence and/or location, it is conceivable that they could be engineered as a multi-epitope vaccine that would be broadly reactive and elicit protective immunity against endemic systemic mycoses. A recombinant chimeric polypeptide antigen (rCpa1) has been genetically engineered encompassing five epitopes derived from *Cp-*specific aspartyl protease (Pep10), a mannosidase (Amn1), and phospholipase B (Plb) antigens (14, 22). The vaccine elicits a mixed Th1 and Th17 immune response and confers protection in C57BL6 and HLA-DR4 mice against multiple isolates of *Cp* and *C. immitis* (23).

We used long “promiscuous” peptides identified in HLA-DR4 mice to survey naïve human subjects for reactive T cell precursors. CD14^+^ monocytes from these subjects loaded with Eng2 homologues or the immunogenic peptides primed and expanded autologous naïve CD4^+^ T cells and triggered the production of multiple cytokines. Interestingly, peptide primed CD4^+^ T cells produced larger amounts of cytokines when recalled with the corresponding peptides than did protein primed T cells. Increased cytokine production is likely due to the fact that peptide-primed T cells were not subject to clonal competition, whereas protein-primed T cells had to compete for antigen and deal with clonal competition. While only peptide 1 for *Cp*-Eng2 was immunogenic in vaccinated HLA-DR4 mice, peptides 1-3 were found to be immunogenic in the MIMIC System assay using naïve T cells. The diversity of HLA haplotypes of the human T cells likely accounted for the recognition and immunogenicity of the additional peptides.

We hypothesized that human infection with these dimorphic fungi elicits adaptive immunity to the Eng2 homologues, given their immunodominance and vaccine effect. Our observation that pulmonary infection of HLA-DR4 mice with *Hc* yeast primed antigen-specific CD4^+^ T cells that produce IFN-γ upon recall *in vitro* with *Hc*-Eng2 supported this hypothesis and prompted us to study Eng2-specific T cells responses in individuals recovered from the respective infections. Such individuals harbored memory T cells that produced cytokines in response to recall *in vitro* with Eng2 homologues indicating that human T cells recognized this antigen during natural infection. These memory T cells were selected by HLA restriction and subjected to antigen and clonal competition. Peptides 1-3 from *Cp*-Eng2 triggered IFN-γ production by memory T cells but the epitope hierarchy was dominated by peptide 3, which was different than the balanced epitope hierarchy we observed with naïve T cells in the MIMIC System. Thus, cytokine production by human memory T cells in response to immunogenic peptides is likely a consequence of several factors, including HLA restriction (e.g. some peptides are not presented by each HLA during infection), immunodominance, precursor frequency and hierarchy of T cell epitopes in response to the fungal infection, and the polyclonal nature of the T cell response.

From this study, we conclude that Eng2 homologues are expressed on the infectious or tissue form of these dimorphic fungi during natural infection and elicit functional memory CD4^+^ T cells with clinical significance. Our findings have implications for strategies to prevent and also reliably diagnose the systemic endemic mycoses of north America, which represent a significant and growing public health problem.

## Author contributions

Designing research studies: MW, BK, GK, CH

Conducting experiments: UO, CLT, LSD, HD, AC, GD

Predicting peptides: JH

Acquiring data: UO, CLT, LSD, HD, AC, GD

Analyzing data: UO, CLT, LSD, HD, AC

Providing reagents: GT

Writing draft of the manuscript: UO, CLT

Editing manuscript: MW, BK

## Acknowledgement

The work was supported by NIH grants BAA-NIAID-DAIT-AI201800007, Contract Number: 75N93019C00064, R01AI93553 (MW), R01AI040996 (BK/MW), R01 AI168370 (BK), U01 AI124299 (BK), R37 AI035681 (BK), R01AI178010 (GSD), R01AI135005 (CYH), U19AI166761 (CYH) and by the Division of Intramural Research of the National Institute of Allergy and Infectious Diseases, NIH (LSD), and the American Heart Association Postdoctoral Fellowship #835129 (LSD). Flow samples were processed at the University of Wisconsin Carbone Cancer Center (UWCCC) Flow Core Facility on a BD LSR Fortessa that was purchased with the NIH shared instrumentation grant 1S100OD018202-01 and University of Wisconsin Carbone Cancer Center Support grant P30 CA014520. Microscopy was performed at the University of Wisconsin–Madison Biochemistry Optical Core (RRID:SCR_023952), which was established with support from the University of Wisconsin–Madison Department of Biochemistry Endowment”

## METHODS

### Sex as a biological variable

Both male and female C57BL/6 and HLA-DR4 mice were used in vaccination experiments and similar outcomes were found. Our human studies included both male and female subjects without explicit analysis dedicated to determination of sex as a biological variable.

### Fungi

Wild-type *Bd* strain ATCC 26199, *Cp* virulent clinical isolate C735 (ATCC 96335), *Hc* G217B and *C. albicans* strain ATCC SC5314 were used for this study. *Bd* 26199 was grown as yeast on Middlebrook 7H10 agar with oleic acid-albumin complex (Sigma) at 39°C. Saprobic phase *Cp* was grown on 2x GYE agar (2% glucose, 1% yeast extract, 1.5% agar) at 30C to produce spores as reported (24). Culture and harvest of *Cp* were done in a biosafety level 3 (BSL3) laboratory located at the University of Wisconsin (UW)-Madison. *Hc* was grown on brain heart infusion agar-blood at 37°C for 15 days. *Candida* yeast was grown on YPD agar at 30°C.

### Mouse strains

Inbred wild-type C57BL/6 mice obtained from Jackson Laboratories were bred at our facility. Male and female mice were 7 to 8 weeks old at the time of these experiments. A breeding colony of HLA-DR4 (DRB1*0401) transgenic mice that express human MHC class II molecules (40)(15) was used in this study. HLA-DR4 mice were engineered from C57BL/6 background and backcrossed to MHC class II-deficient mice lacking IA and IE alleles to eliminate production of endogenous murine MHC class II. Mice were housed and cared for in a specific-pathogen-free environment in our animal facility as per guidelines of the University of Wisconsin Animal Care Committee, who approved all aspects of this work. Mice were transported to the animal BSL3 (ABSL3) laboratory for challenge with live *Cp* spores.

### Immunoinformatics and epitope mapping

Immunogenic peptides were predicted using Eigenbio immunoinformatic platform described elsewhere (25, 26). Briefly, the mean and standard deviation (SD) of the natural log of IC50 MHC II allele binding for each sequential 15-amino-acid peptide in the protein were predicted by artificial neural network ensembles using algorithms based on vectors derived from the principal components of the physical and chemical characteristics of each amino acid. Mean predicted binding was standardized to a zero mean unit variance (normal) distribution within the protein to provide a relative competitive index of predicted binding for each peptide in the protein. This places binding predictions of all MHC alleles on the same scale. This metric is expressed in SD units relative to the mean for that protein. Comparison with other prediction systems indicates that a predicted binding affinity of less than -1 SD units below the mean is a probable epitope (27). Predicted peptides were synthesized at >75% purity by GenScript and provided as lyophilized material in measured amounts of peptide. Each peptide was analyzed by GenScript for purity using high-performance liquid chromatography (HPLC), mass spectrometry, and nitrogen analysis. Depending on their solubility, peptides were dissolved in water or 100% dimethylsulfoxide (DMSO). Stock solutions of each peptide were adjusted to 10mM and stored at -80°C.

### Generation and purification of recombinant Eng2 homologues

*Cp*-Eng2 and *Hc*-Eng2 were cloned and expressed in *P. pastoris* and *E. coli* using standard recombinant techniques and have been described for Bl-Eng2 (28). Recombinant proteins were purified using Ni-NTA agarose (Qiagen) according to the manufacturer’s protocol and dialyzed against PBS. Purity of recombinant proteins was assessed by SDS-PAGE and silver staining.

### Generation of rabbit anti-Eng2 antibody and for surface staining of dimorphic fungi

Rabbit polyclonal antibody was generated by Envigo Research and Model Service, Madison, WI. Briefly, rabbits were vaccinated thrice with 1mg of recombinant of each Eng2 homologue and serum harvested on day 112 after first vaccination.

Yeast or spherules were washed and fixed in formaldehyde 4% for 15 minutes at room temperature, washed, resuspended in blocking buffer containing 1% BSA in PBS + 0.1% Tween 20 for 15 minutes. After washing, fungi were stained with the corresponding polyclonal anti-Eng2 rabbit antibodies at a 1:800 dilution and incubated at 4°C overnight. After washing, fungi were stained with 5 µg/ml of a secondary Donkey anti-Rabbit IgG antibody conjugated with Alexa Fluor™ 647 (Invitrogen) for two hours. After washing, fungi were stained with 10 µg/ml of Calcofluor White Blue (Sigma) for 20 minutes and washed again. Samples were resuspended in mounting medium (VECTASHIELD® Vibrance™ Antifade Mounting Medium [VectorLabs], transferred to microscope slides, covered with a coverslip [Circular Coverglass 12mm (Electron Microscopy Sciences) and stored at 4°C. A Nikon A1R confocal microscope was used to capture the images of the slides.

### Vaccination and fungal infection

Ten micrograms of *Bl*-Eng2, 20μg *Cp*-Eng2 or 30μg *Hc*-Eng2 or equimolar amounts of synthetic peptides were loaded into GCPs. For vaccination with peptide pools, individual peptides were resuspended in DMSO and equal amounts of each peptide were pooled and mixed with 50% CAF01 (Statens Serum Institute, Copenhagen, Denmark) or 10% adjuplex (Empirion, LLC, Columbus OH). Mice were vaccinated subcutaneously (SC) with Eng2 full length proteins three times, two weeks apart as described (29) with antigen loaded into glucan chitin particles (GCPs); controls in these experiments were GCPs loaded with mouse serum albumin (MSA). In limited experiments, Freund’s adjuvant was used for vaccination. Two weeks after the vaccine boost, mice were challenged intratracheally (i.t.) with 2×10^4^ wild type *Bd* yeast, 100-150 *Cp* spores or 10^6^-10^7^ *Hc* yeast in 30μl PBS. At day 4-6 post-infection, lung T cell responses were analyzed and lung CFU counted. Two weeks post infection, when control or unvaccinated mice were moribund, fungal burden was determined by plating lung CFU.

### Generation of MHCII tetramer

Tetramers for detection of Eng2-specific CD4^+^ T cells in C57BL6 were generated at the NIH Tetramer core facility at the Emory University in Atlanta, GA. A tetramer that recognizes *Cp*-Eng2-specific CD4^+^ T cells in HLA-DR4 mice was generated by ProImmune, Oxford, England.

### Human donors and PBMC isolation

For MIMIC® assays, PBMCs were obtained from healthy donors who enrolled in a Sanofi VaxDesign apheresis study (protocol CRRI 0906009) after informed consent. Leukocytes were enriched by centrifugation over a Ficoll-histopaque PLUS (General Electric Healthcare, Piscataway, NJ) density gradient (33), collected at the gradient interface, washed, and cryopreserved in Iscove’s Modified Dulbecco’s Medium (Lonza, Walkersville, MD) containing autologous serum and 10% DMSO (Sigma-Aldrich, St. Louis, MO). Blood products were negative for blood-borne pathogens as detected by standard assays. For bloods in patients recovered from infection, cases of blastomycosis and coccdioidomycosis were confirmed by culture or histopathology. For histoplasmosis, we received deidentified blood from the Hoxworth blood bank, University of Cincinnati College of Medicine.

### Modular Immune in vitro Construct (MIMIC) System CD4^+^ T Cell LTE Assay

Sanofi’s proprietary MIMIC System was used to assay whether naïve human CD4^+^ T cells recognize peptides in Eng2 homologues. Human donors were selected based on their HLA-DRB1 haplotypes to achieve a diverse sampling of more commonly expressed HLA-types. CD14^+^ cells and CD4^+^ T cells were purified from PBMCs of donors using CD14^+^ positive and CD4 negative selection isolation kits (Stem Cell Technologies, Vancouver, Canada). CD14^+^ cells were treated with recombinant IL-4 and GM-CSF to promote DC differentiation. One week post-isolation, these cytokine-derived dendritic cells (CDDCs) were loaded with antigens at doses ranging from 1-5 mg/ml. A vehicle control was included to assess donor background responses. Antigen-loaded DCs were co-cultured with purified donor-matched CD4^+^ T cells for 14 days, as part of the primary stimulation phase. Unpulsed DCs, or those pulsed with irrelevant antigen served as negative controls.

After two weeks, co-cultured cells were harvested and restimulated with fresh CDDC loaded with the same protein or peptide or related antigen. Cells were then assayed for CD154^+^ expression (activation) and intracellular cytokine. To assay intracellular cytokines, cultures received Brefeldin A for 5-7 hours of restimulation. After restimulation, cells were fixed, permeabilized, and stained with a panel of antibodies to detect intracellular IFN-γ, TNF-α, and IL-2 (demark Th1 cells), IL-4, IL-5, and IL-10 (Th2 cells) and IL-17 (Th17 cells). Data were analyzed with FlowJo, Graphpad Prism, Microsoft Excel, and Spice (v5.3).

### ELISA

Cytokine concentrations in cell culture supernatants were determined by IFN-γ and IL-17A Duoset ELISA kits (R&D Systems, Minneapolis, MN) according to manufacturer’s instruction.

### Tetramer pulldown

Miltenyi LS columns on a quadroMACS magnet were used to enrich tetramer^+^ cells from secondary lymphoid organs (SLO). Spleen and draining lymph nodes were harvested and mashed through 40 μm filters. Red blood cells were lysed with ACK buffer. Samples were washed with RPMI and resuspended in cold sorter buffer (PBS with 2% FBS) to a volume twice the size of the pellet. Two microliters of Fc block was added to each sample and incubated for 5 min before adding tetramer [5–25 nM]. Tetramer stain was done for 1 h at room temperature in the dark. Samples were then washed and kept on ice. One hundred microliters of Miltenyi anti-PE microbeads (Miltenyi 130–048–801) was added to each sample and incubated for 30min on ice, washed and resuspended in 3 mL of sorter buffer. LS columns were pre-wet and samples filtered through 40 μm filters into fresh columns. Columns were washed with cold sorter buffer thrice before eluting bound fractions. Fractions were stained with Invitrogen’s LIVE/ DEAD^TM^ stain and surface markers. AccuCheck Counting Beads (50μl) were added to each sample to determine total number of tetramer positive cells.

### T-cell stimulation, and flow cytometry

Lungs were dissociated in Miltenyi MACS tubes and digested with collagenase (1 mg/mL) and DNase (1 μg/mL) for 25 min at 37°C. Digested lungs were resuspended in 5 mL of 40% percoll; 3mL of 66% percoll was underlaid (GE healthcare, cat# 17–0891–01). Samples were spun for 20 min at 2,000 rpm at room temperature. Lymphocytes in the buffy coat were harvested and resuspended in RPMI (10% FBS, 1% penicillin and streptomycin). For T-cell stimulation *ex vivo*, cells were incubated at 37°C for 5h with 5 μM peptide and 1μg anti-mouse CD28 (BD 553294). After 1h, BD GolgiStopTM (BD, cat# 554724) was added to samples. FACS samples were stained with Invitrogen’s LIVE/DEAD^TM^ stain and Fc block for 10 min at room temperature. Cells were stained with tetramer for 1h at room temperature, or for surface antigens or intracellular targets for 20 min at 4°C. Transcription factors were stained on primed, resting cells using the Foxp3 Transcription Factor Staining kit (ebioscience cat# 00– 5523–00). All panels included a dump channel (Dump markers:CD11b, CD11c, NK1.1, CD19 B220, CD8). 50 μl AccuCheck Counting Beads (Invitrogen PCB100) were added to samples to determine absolute cell counts. Samples were acquired on a LSR Fortessa.

### Ethics Statement

Research was performed in accordance with approvals from the Human Research Ethics Committee of the UW-Madison. Participants provided written informed consent. Animal studies adhered to protocol M005891 approved by the IACUC of UW-Madison. Animal studies were compliant with provisions established by the Animal Welfare Act and the Public Health Services (PHS) Policy on the Humane Care and Use of Laboratory Animals.

### Statistical analyses

A Student t test was used to analyze the differences between two treatment groups for cytokine ELISAs, ELISPOT assays, cytokine concentrations, calculations of numbers of lung-infiltrating immune cells, and percentages of specific-cytokine-producing T cells. Differences in fungal burden (expressed as CFU) between two groups were analyzed by the Mann-Whitney U test for ranking data. For comparison of fungal burden among three or more groups of mice, the Kruskal-Wallis test, a nonparametric ranking method, was used. Survival data were examined by the Kaplan-Meier test using log rank analysis to compare survival plots as reported previously (6). A p value of ≤0.05 was considered statistically significant. Comparisons in many experiments yielded p values at or below the value of p = 0.05, however we consistently used only one asterisk throughout to denote any statistically significant difference regardless of the exact value below 0.05.

## SUPPLEMENTAL FIGURE LEGENDS

**Supplementary Figure 1 (related to Figure 2):**
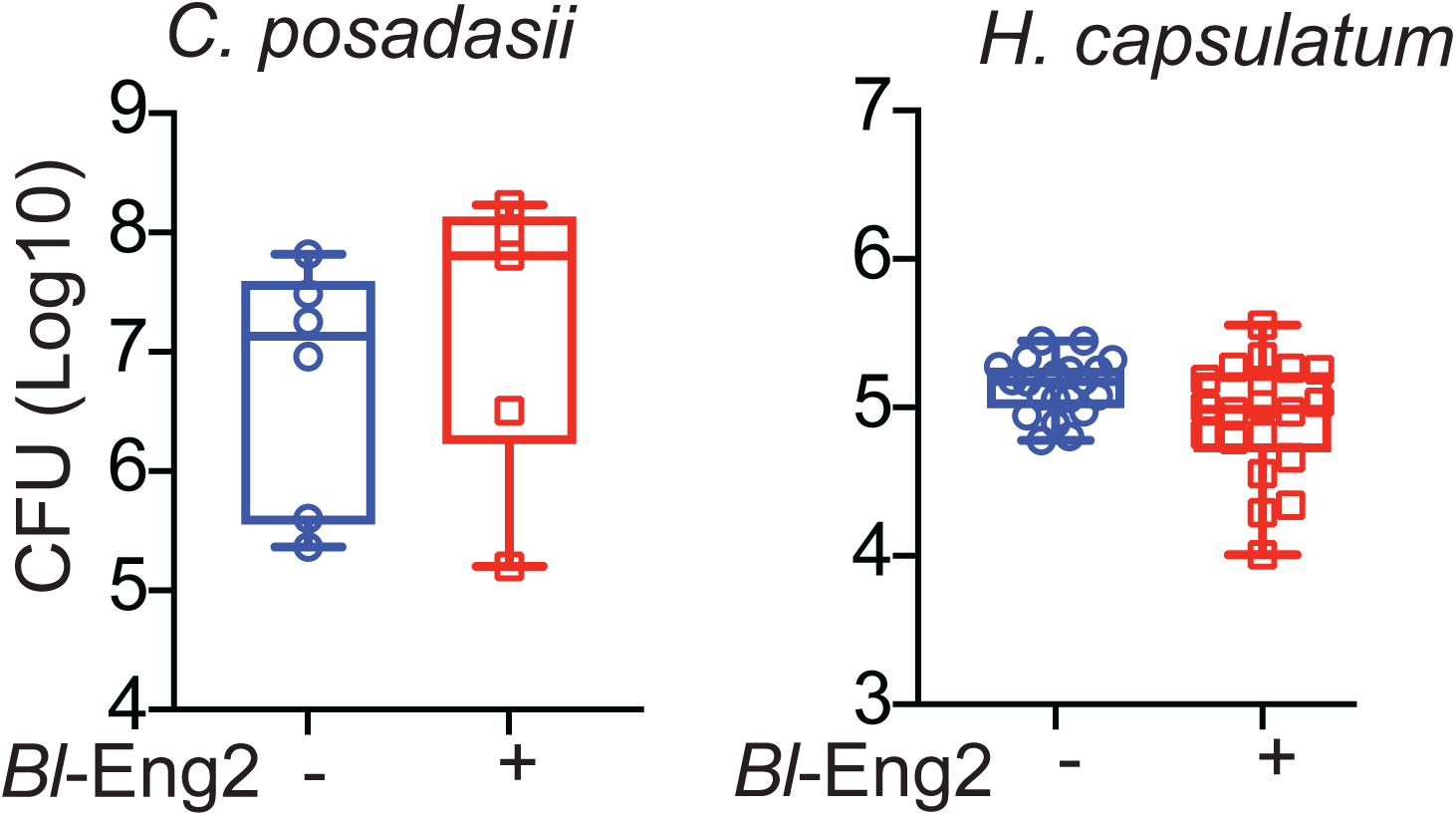
Protective eaicacy of Bl-Eng2 against infection with *C. posadasii* and *H. capsulatum*. C57B/6 mice were vaccinated with *Bl*-Eng2, challenged and analyzed for lung CFU as described in Methods. Vaccinated mice were challenged with *Cp spores* or *Hc* yeast i.t. as indicated. Data shown are from n=6-10 mice/group. A representative experiment is shown of two performed. CFU are expressed as Log_10_ plotted with geometric mean ± geometric SD. *p<0.05, two tailed Mann-Whitney U t-test.

**Supplementary Figure 2 (related to Figure 3).**
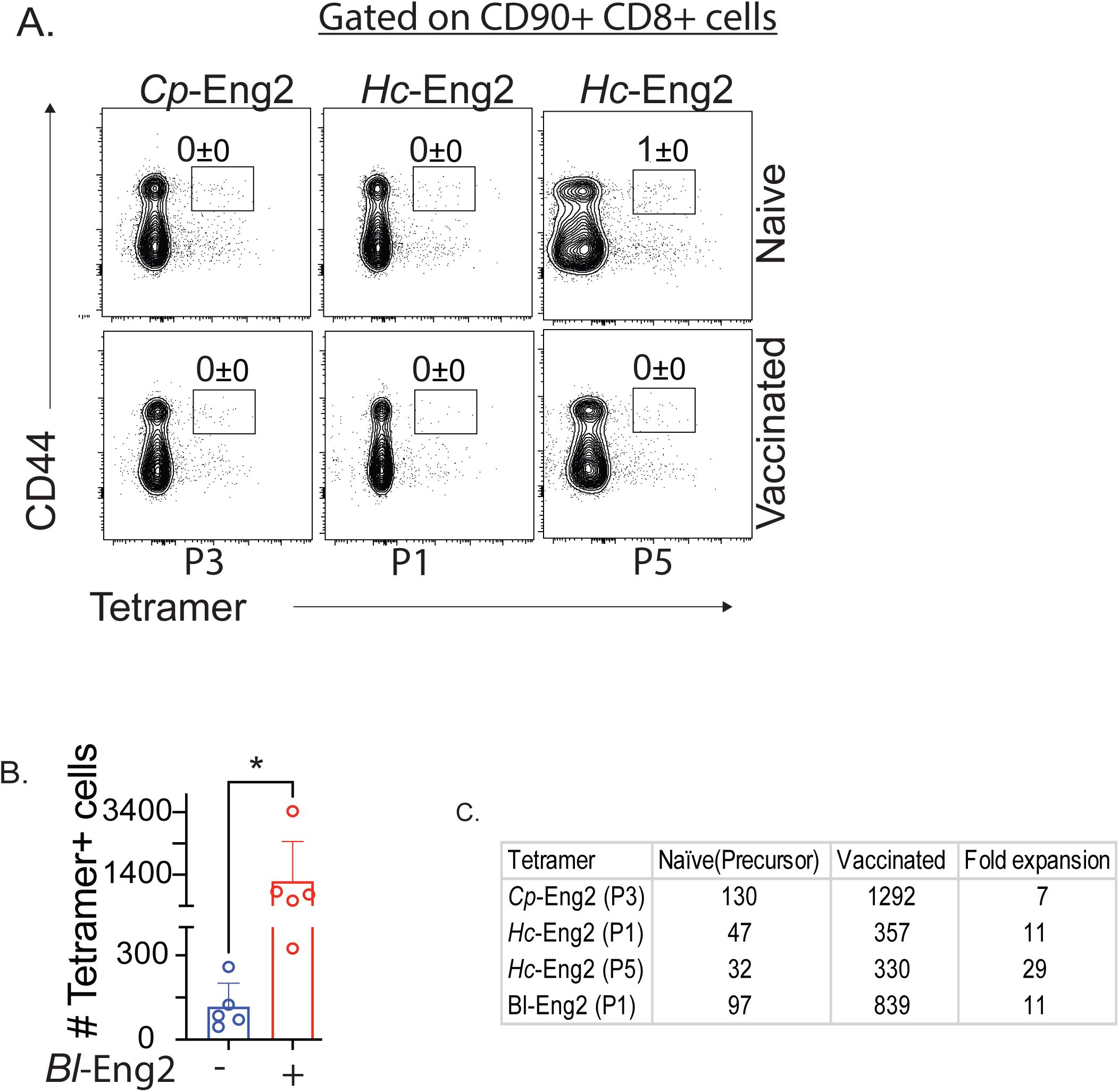
Analysis of Eng2-specific tetramers. **(A)** CD8^+^ T cells from vaccinated C57BL/6 mice were stained with MHC class II tetramers containing peptides P3 (*Cp*-Eng2), P1 or P5 (*Hc*-Eng2) and analyzed by flow cytometry. Dot plots are concatenated data from 5 mice and show the percentage of CD8^+^ T cells that stained with the tetramers. The data are representative of 2 independent experiments. Data are represented as mean± SEM. Statistical analysis was using Mann-Whitney U t-test *p=<0.05. (**B)** The number of CD4^+^ T cells from the spleen and draining lymph nodes of *Bl*-Eng2 vaccinated mice that stain with the Bl-Eng2-specific tetramer prior to challenge. (**C)** Tetramers specific to the corresponding homologues of Eng2 were used to stain CD4^+^ T cells from the spleen and lymph nodes of naïve and vaccinated mice. Data shown are from n=5 mice/group.

**Supplementary Figure 3 (related to Figure 6).**
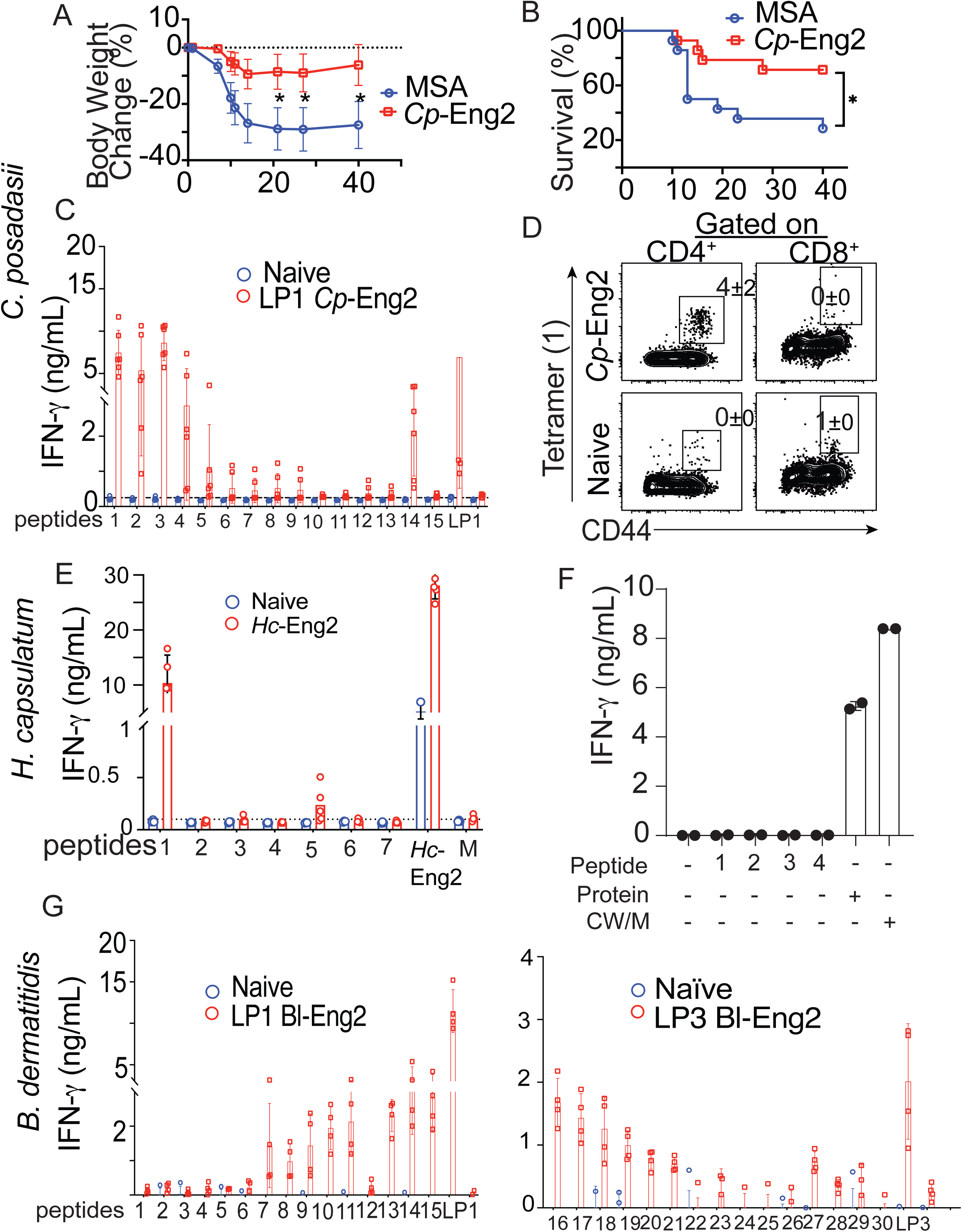
Resistance to *Cp* and epitope mapping in vaccinated HLA-DR4 mice. Resistance to infection in vaccinated mice after challenge as measured by weight change **(A)** and (**B)** survival. **(C)** Identification of immunogenic 15mer from long peptide 1 (25-30 amino acids) *Cp*-Eng2. **(D)** Validation of tetramer for detection of *Cp*-Eng2-specific T cells. **(E)** *Hc*-Eng2 protein was mapped for the immunogenic 15-mer (note that peptide number 1 is contained within the sequence of long peptide 3), **(F)** HLA-DR4 mice were infected intranasally with *H. capsulatum* G217B. One month later splenocytes were stimulated with *Hc*-Eng2 peptides, protein or CW/M extract and IFN-γ measured in the cell culture supernatants. **(G)** *Bd*-P1 and *Bd*-P3 were mapped for their immunogenic 15-mers.

**Supplementary Figure 4 (related to Figure 6).**
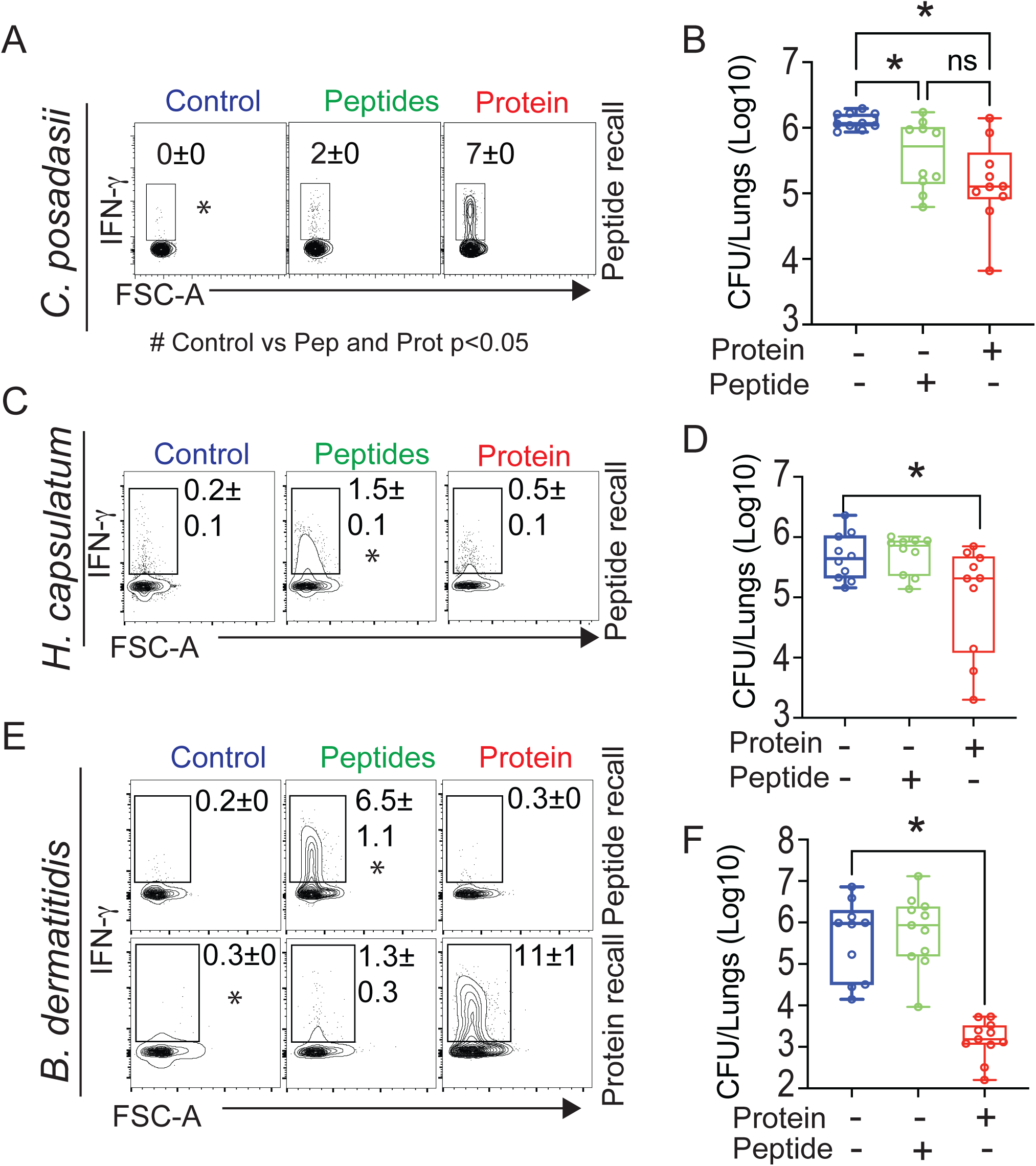
Vaccine protection conferred by immunodominant peptide vs. full-length protein homologue in humanized DR4 mice. Mice were vaccinated with 20μg *Cp*-Eng2, 30 μg *Hc*-Eng2 and 10μg *Bl*-Eng2 and equimolar amount of immunodominant peptides: LP1 and peptides 1, 2, 3, 4 from *Cp*-Eng2, LP3 and peptides 1 and 5 from *Hc*-Eng2 and LP1 and peptides 13, 14, 15 and LP3 and peptides 16, 17 and 18 from Bl-Eng2) and challenged with a lethal dose of each organism i.t. as described in methods. Single cells from the lungs were stimulated ex-vivo with peptides (as used for vaccination) or protein (as indicated) and stained for intracellular cytokine stimulation at day 6, 5, and 4 post challenge with *Cp, Hc* and *Bd* respectively. **(A, C & E)** The percentage of cytokine-producing cells in the lung in response to *Cp, Hc* and *Bd* challenge respectively. **(B, D & F)** Resistance to infection as measured by lung CFU at endpoint when the control group was moribund (n=10mice/group). *p<0.05 vs. all other groups.

